# Efficient and Explainable Deep Neural Networks for Airway Symptom Detection in Support of Wearable Health Technology

**DOI:** 10.1101/2021.12.30.474418

**Authors:** René Groh, Zhengdong Lei, Lisa Martignetti, Nicole Y. K. Li-Jessen, Andreas M. Kist

## Abstract

Mobile health wearables are often embedded with small processors for signal acquisition and analysis. These embedded wearable systems are, however, limited with low available memory and computational power. Advances in machine learning, especially deep neural networks (DNNs), have been adopted for efficient and intelligent applications to overcome constrained computational environments. In this study, evolutionary optimized DNNs were analyzed to classify three common airway-related symptoms, namely coughs, throat clears and dry swallows. As opposed to typical microphone-acoustic signals, mechanoacoustic data signals, which did not contain identifiable speech information for better privacy protection, were acquired from laboratory-generated and publicly available datasets. The optimized DNNs had a low footprint of less than 150 kB and predicted airway symptoms of interests with 83.7% accuracy on unseen data. By performing explainable AI techniques, namely occlusion experiments and class activation maps, mel-frequency bands up to 8,000 Hz were found as the most important feature for the classification. We further found that DNN decisions were consistently relying on these specific features, fostering trust and transparency of proposed DNNs. Our proposed efficient and explainable DNN is expected to support edge computing on mechano-acoustic sensing wearables for remote, longterm monitoring of airway symptoms.

## Introduction

Wearable device technology is widely adopted in the healthcare community. For complex disease diagnosis and monitoring, multiple physiological signals are continuously streamed to a wearable device and multiple decisions need to be intelligently made within a short time window. The integration of artificial intelligence (AI) into smart wearable devices is particularly needed for effective and accurate processing of health data at the point of care. Most wearable devices are embedded with a sensor, a microprocessor and a limited memory flash to keep the system small and lightweight. However, such constrained computational environments make the deployment of advanced AI techniques very challenging (1).

Cough and other audible sounds (e.g., wheezing, deviated voice quality etc.) have been used as digital audio biomarkers for early disease detection or predicting acute exacerbations in airway diseases such as asthma, chronic obstructive pulmonary diseases (COPD) and COVID-19 (2–4). Most wearable health devices for airway diseases are built on audio sensing technology to detect aforesaid airway symptoms with an embedded microphone (5). These acoustic microphones are often omnidirectional and capture both, a speaker’s voice and surrounding sounds. Wearing a constantly recording microphone creates inevitable personal privacy concerns.

As a promising alternative, mechano-acoustic sensing devices, such as those of neck surface accelerometers (NSAs), are noise resistant and equally capable of generating airway health-related information (6–9). An NSA device detects and transfers mechanical vibrations from the neck skin to electrical voltage signals. The sensor captures negligible vocal tract resonance information during phonation, which preserves a person’s speech privacy (10, 11). The sensor is also insensitive to air-borne acoustic waves (12), which ensures highquality data acquisition due to its intrinsic anti-interference against background noise. On the other hand, the attenuation of frequency information in NSA signals may make the AI classification tasks more challenging compared to that of microphone-acoustic signals.

AI and related deep learning technologies have been shown to accelerate the time course and improve the quality of disease diagnosis and treatment monitoring (13, 14). Recently, sophisticated AI methods have been adopted for classifying airway-related symptoms such as cough (15, 16) and deviated voice quality (17) in various clinical populations. Lean models have been proposed for the detection of cough in patients suffering from chronic cough, COPD, asthma and lung cancer (5). Cough detection is also helpful in predicting COVID-19 infection (4). However, these deep neural networks (DNNs) have barely been optimized for wearable devices. Further, not many algorithms are capable of multiclass classification in detecting more than one airway symptom

(18) Also, given the black-box-character of AI algorithms, explainable AI has been advocated to increase trust, especially in the development of health wearable devices (19, 20). In this work, we aimed to optimize multiple neural network topologies using evolutionary algorithms to allow explainable, personalized airway symptom detection on a wearable device (Fig. 1 for study overview). In this work, research questions were:

**Fig. 1.**
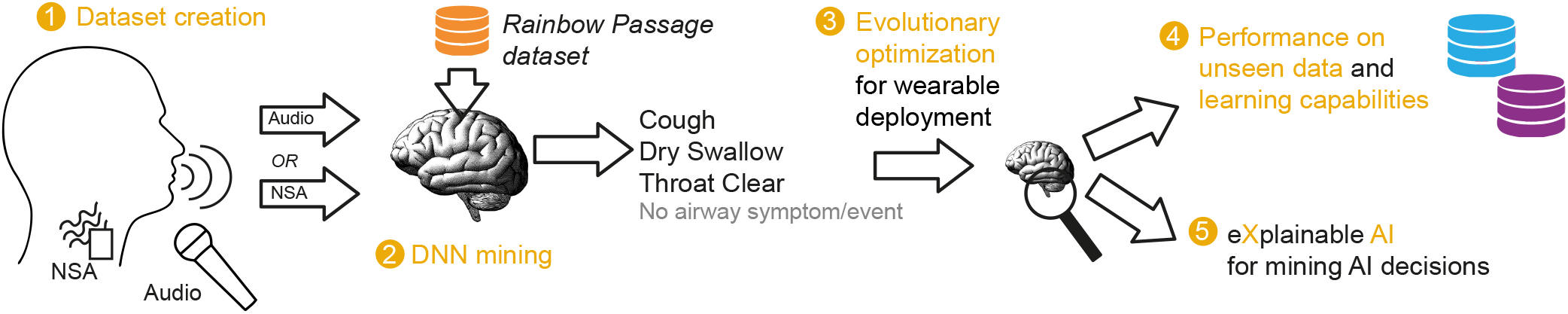
Overview of this study as flow diagram. Orange circles indicate milestones in the project, where the milestones 1-5 are reflected by the Figures 2-6 in this study.

- Would NSA signals be on par with audio signals in terms of classification accuracy?
- Which AI technologies would suit for classifying airway symptoms?
- Would evolutionary algorithms be capable of optimizing DNN topologies to gain neural networks for the deployment on wearable devices?
- Would the optimized DNNs be able to cope with new, unseen datasets?
- Would the optimized DNNs rely on specific features, i.e. frequencies of NSA signals in airway symptom classification?

## Methods

### Datasets

Three individual datasets, which contained airway symptoms of interests, were curated from laboratory-generated or public sources. These datasets were from: (1) a study of reading a standard passage scripted with airway symptom productions (Rainbow Passage dataset), (2) a published study of vocal loading tests (Vocal Stress dataset) (9) and (3) a crowdsourcing COVID-19 cough sound project (COUGHVID dataset) (21).

#### Rainbow Passage dataset

This human study was approved by McGill University Research Ethics Office (A11-B62-19A). All participants of this study gave their informed, written consent. Six female adult participants, who were vocally healthy with age ranging between 20 and 35, were recruited to this experiment. Both audio (ICD-UX565F, Sony Inc., Japan) and NSA data were recorded simultaneously (Fig. 2A) in a sound-proof booth. Participants were first prompted to produce isolated cough, throat clear and dry swallow sounds. Participants were then asked to read the Rainbow Passage, which was scripted with the three airway symptoms interspersed throughout, using their conversational pitch and loudness. They read this script three times in a row (Fig. 2B, Supplementary Material).

**Fig. 2.**
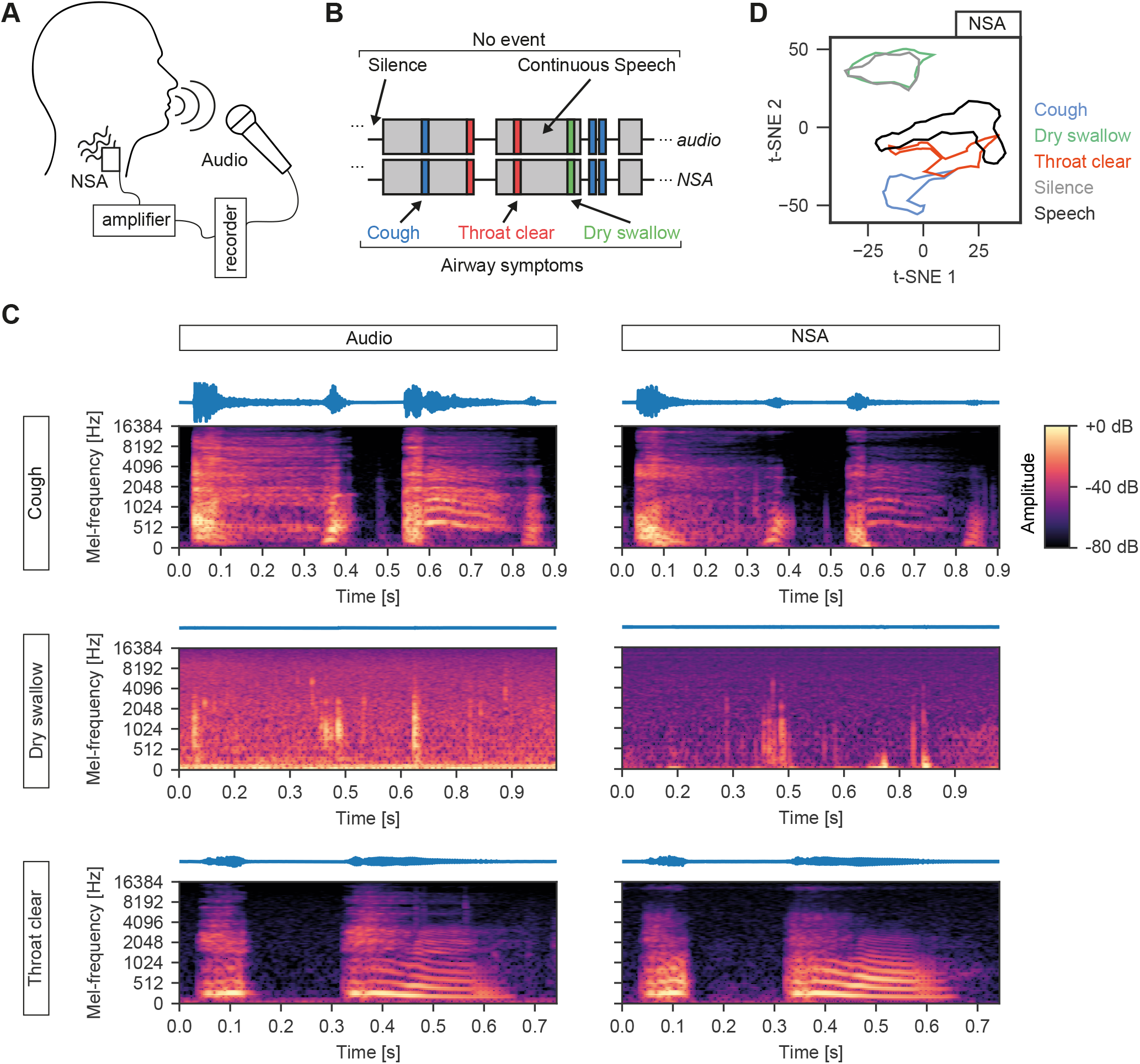
Airway symptoms in NSA signals. **A** Recording condition of the Rainbow Passage dataset. **B** Schematic of the annotated paired audio and NSA data. Silence and speech are labelled both as “no event”, whereas cough, throat clear and dry swallow are distinct categories. **C** Representative log-mel-spectrograms of paired audio and NSA signals for each airway symptom. **D** t-SNE representation of all categories described in panel B.

The main unit of the NSA was a printed circuit board embedded with a one-axis accelerometer (BU-27135, Knowles Inc., IL, USA) and a custom amplification module to preprocess and transmit the signal to a recording device. A total of 294 coughs, 287 dry swallows and 382 throat clears were obtained in this dataset. Fig. 2C shows representative examples (paired audio/NSA signals and the corresponding logmel-spectrograms) for the three symptom classes. Data were annotated by two experts who had more than five years of clinical voice evaluation experience, using a custom graphical user interface written in Python.

#### Vocal stress task dataset

In addition to the Rainbow Passage dataset, we sought to obtain data samples of airway symptoms that were elicited in a relatively natural setting. Our published data set, in which the airway symptoms were produced spontaneously by speakers during a vocal stress task, was selected for this study (9). In brief, nine female adult participants were asked to read parts of the novel “Harry Potter and the Sorcerer’s Stone” (22) for up to 3 hours. Both audio and NSA signals were obtained in a sound-booth environment using the same devices as those of aforesaid Rainbow Passage dataset experiment. A total of 19 coughs, 258 dry swallows and 11 throat clears were annotated in this dataset.

#### COUGHVID dataset

Cough is one most common symptoms in airway disease diagnosis and monitoring. To further evaluate our AI algorithm, a highly heterogeneous dataset of coughs containing more than 20,000 recordings were collected from the COUGHVID crowdsourcing dataset (21). The predictions of the classifiers were already stored in the original COUGHVID data files by (21). We thus preselected the cough admissions with more than 98% classifier probability. Given that non-cough parts were also contained in the recordings, we computed the rolling standard deviation with a window size of 5,000 sampling points and an energy threshold of 8,000 to determine the onset of the cough event. As a result, a total of 3,388 cough events were obtained for the evaluation of our AI algorithms in this study. Of note, as these cough sounds were microphone audio signals, an autoencoder DNN architecture was applied to convert the audio samples to NSA space (Supplementary Fig. 5).

### Data preprocessing

During preprocessing, both audio and NSA data were divided into 500 ms frames. For each frame, the mel-spectrogram was calculated using 64 mel-frequency-bands, an FFT window length of 1024, a hop length of 64, an upper frequency bound of 16384 Hz and the HTK-formula (23) for conversion from Hertz to mel. The advantage of mel-spectrograms is that the center frequency and bandwidth of the chosen triangular filters roughly match the auditory critical band filters (24). Using the Python package librosa (25), each 500 ms frame resulted in a mel-spectrogram with 64 frequency points and 345 time frames.

Other preprocessing steps included calculating the logmel-spectrogram, scaling the values in the range of -1 to 1 (min-max normalization), flipping the spectrogram such that lower frequencies were at the bottom of the spectrogram, and resizing the spectrogram to 64 *×*64 data points, which we further used as an image-like object in pixels. Finally, a class label was assigned to each of the log-mel-spectrograms. If more than 70% of the 500 ms window belonged to an annotated event, the log-mel-spectrogram was labeled accordingly, i.e., Cough, Dry swallow, Throat clear or No event. These log-mel-spectrograms and their associated labels were treated as inputs and outputs, respectively, to various DNN architectures for the classification of airway symptoms.

### Data visualization

The t-distributed stochastic neighbour embedding (t-SNE) dimensionality reduction technique (26) was used to visualize the relationship of the three airway symptoms. Input to the t-SNE algorithm were log-mel-spectrogram-derived features, including the mean, min, max, median, mode and standard deviation of each coefficient of one mel-frequency band. These extracted features were then projected onto two t-SNE dimensions. The t-SNE algorithm was implemented using the scikit-learn library (27) with the perplexity set as 40, the learning rate as 30, and the number of iterations as 1500. We computed the alpha shape of each class and reported individual clusters to illustrate possible overlapping of multiple classes.

### Network architectures and training

We evaluated five state-of-the-art network architectures in the classification of cough, dry swallow, throat clearing, and no event (Fig. 3A). We focused on convolutional neural networks (CNNs) and recurrent neural networks (RNNs), specifically, the ResNet architecture (ResNet18 and ResNet34 (28)), EfficientNetB0 (29), the MobileNetV2 (30), an Encoder-Decoder-RNN (31), and a vanilla RNN that we developed specifically for this study. The vanilla RNN consisted of three Long Short-Term Memory (LSTM) layers (32) of 128, 64 and 32 cells, respectively, followed by a fully connected layer with softmax activation function.

**Fig. 3.**
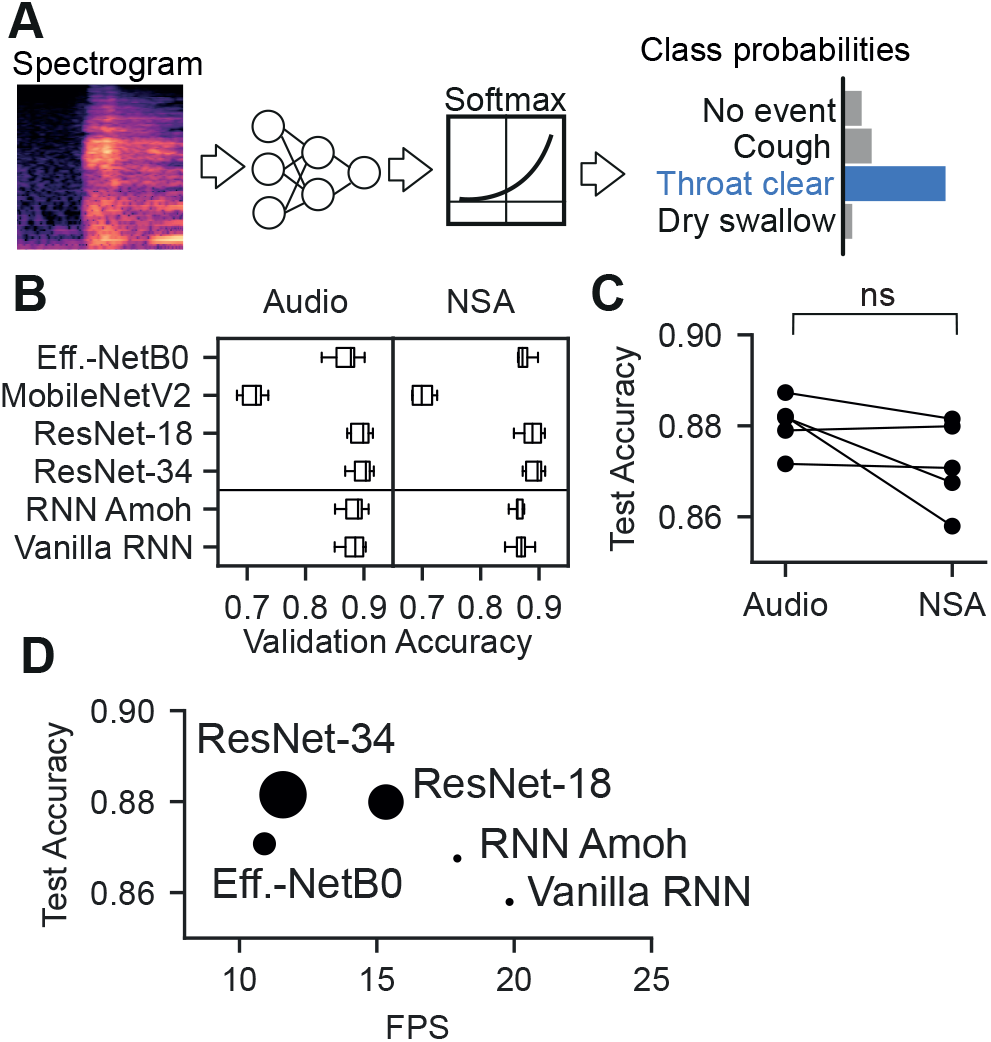
Training of six CNN and RNN architectures for classifying airway symptoms. **A** Overview of the classification pipeline. Log-mel-spectrograms obtained as pre-processing step are the input of neural network architectures for predicting airway symptoms after softmax activation. **B** Validation accuracy for different training/validation splits of the Rainbow Passage dataset for audio and NSA. **C** Median test accuracy from all test accuracies determined by cross-validation. **D** FPS during model inference and median test accuracy from cross-validation for NSA data. The size of the points correlates with the number of trainable parameters of each architecture.

All experiments were implemented using Google TensorFlow (version 2.5.0 with keras API) on an NVIDIA GeForce RTX 3090 GPU and an Intel Core i9-10900X CPU. During network training, the Adam optimizer (33) was used to minimize the categorical cross-entropy loss. The learning rate was 10^*−*4^ with an exponential decay over time. Since the entire dataset had an imbalance of events and non-events, scikit-learn was used to calculate class-dependent weights for model training (27). Models were trained on the whole Rainbow Passage dataset and optimized with the Vocal Stress and COUGHVID datasets. As the Rainbow Passage dataset was generated from six speakers only, a six-fold cross-validation approach was used in the scheme of four speaker dataset for model training, one speaker dataset for model validation and one speaker dataset for model testing.

#### Domain adaptation

COUGHVID crowdsourcing dataset was used to further evaluate the performance of our network architectures in handling complex and heterogeneous data. Since the recordings are microphone audio samples, each COUGHVID sound sample were converted to NSA space for our application. We utilized a U-Net (34) autoencoder that was already trained with the paired audio and NSA signals from the Rainbow Passage dataset. The architecture consisted of four encoder and four decoder layers with 8, 16, 32 and 64 convolutional filters at each depth layer that were connected via skip connections. During the training, mean squared errors were minimized using the Adam optimizer with an exponentially decaying learning rate. We used the tanh-activation function in the output layer to ensure that the output log-mel-spectrograms would contain values in the range from -1 to 1 as noted in the Data Preprocessing section.

### Evolutionary optimization

To find a small and efficient CNN architecture, we utilized an evolutionary algorithm (35). Table 1 shows the gene pool used to determine the neural network topology. The algorithm was allowed to evolve for 20 generations with a population size of 50 in each generation. The individuals of the first generation were created randomly. After each generation, 15 models with the highest fitness scores were selected and used for breeding the next generation’s population. We further employed a mutation rate of 10% during breeding. The fitness for each architecture was calculated using the validation accuracy and the inference time and was defined as follows:

**Table 1.** Overview of the used gene pool in evolutionary optimization.

| Parameter | Set of possible values |
| --- | --- |
| Number of convolutional layers | [1, 2, 3, 4, 5] |
| Number of convolutional filters | [8, 16, 32, 64, 128] |
| Convolutional filter size | [3] |
| Max pooling layer | [True, False] |
| Residual Connections | [True, False] |
| Batch Normalization layer | [True, False] |
| Activation function | [ReLU, ReLU6, LeakyReLU] |

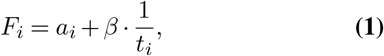

with *F* as fitness, *a* the validation accuracy, *β* the inference time weight and *t*_*i*_ the inference time in seconds of a single frame for each architecture *i*. For objective 1 (O1), we set *β* = 0 to evolve only based on accuracy. For objective 2 (O2), we set *β* = 0.05 to balance accuracy and time-dependence. Each architecture was trained for 12 epochs.

### Microcontroller deployment

To evaluate the scalability of our DNNs, we converted evolutionary optimized models to TensorFlow Lite according to standard procedures. We deployed the converted model to a development board (EdgeBadge, Adafruit Industries) as C array and measured the inference time per single forward pass. An average of 100 single forward passes were reported herein.

### Class activation maps and occlusion experiments for explainable AI

Class activation maps (CAMs) (36, 37) and occlusion experiments (38) were employed to explain neural network decisions. We created the CAMs for each logmel-spectrogram of the test split of the Rainbow Passage dataset. We calculated the weighted sum of each output of the last convolutional layer as described in (36). The class weights of the last network layer (fully connected layer) were used for weighting purposes. Further, we averaged all resulting CAMs of each input log-mel-spectrogram to determine which mel-frequency bands would be of high importance for classification.

The occlusion experiments were performed using a sliding window of size 16*×* 16 pixels and a stride of four pixels. The values in the windowed regions were set to -1 to hide the corresponding information. We then used our trained neural networks for inference to obtain and store the corresponding prediction probabilities for each occluded log-mel spectrogram. Due to overlapping windows, we averaged the pixel values gained from the multiple predictions for reporting purposes.

## Results

### Detectable airway symptoms from NSA signals

Labels were created for “no event” (continuous speech and silence) and “event” (cough, throat clear and dry swallow) of the Rainbow Passage dataset during the expert annotation task (Fig. 2B). Representative audio and NSA data pairs for the three airway symptoms are shown in Fig. 2C. We found that audio and NSA data shared qualitative similarities in the low frequency bands. Given the low-pass filter quality of the NSA, higher frequencies were less present as expected. Distinctive clusters were noted in cough, throat clear and speech (Fig. 2D) under t-SNE representation. Whereas the clusters of dry swallow and silence were highly overlapping, which might lead to difficulty in detecting dry swallow events. Based on these results, airway symptom information were reliably preserved and detected in NSA signals.

### Multi-class classification of airway symptoms with DNNs

Two major DNN technologies, namely CNNs and RNNs, were evaluated for their suitability of airway symptom detection. A library of standardized log-mel-spectrograms was generated from the annotated Rainbow Passage dataset to train, validate and test each DNN by forcing the network to choose one of the four classes (Fig. 3A). The examined CNNs and RNNs were found to operate within similar validation accuracy range and were largely independent of the recording modality (Fig. 3B, Supplementary Fig. 1 and 2). MobileNetV2 was the only architecture that notably underperformed compared to other architectures (Fig. 3B).

We next determined the test accuracy for each architecture. The test accuracy for each architecture in airway symptom prediction was slightly worse with NSA signals but not statistically significant compared to audio signals (Wilcoxon test of paired samples, p=0.19) (Fig. 3C). CNN-based models were also found to be more accurate than RNN-based models. However, CNN-based models were in general slightly slower in terms of fps (Fig. 3D). The distribution of all test accuracies across all cross-validations can be found in detail in Supplementary Fig. 3.

Subsequently, we evaluated if the accuracy of a CNN-based model could be traded for inference speed. As baseline, we chose the ResNet-18 architecture, as it provided a high test accuracy of 88.0% as well as 15.3 fps in classifying airway symptoms, and was already a smaller variant of the ResNet-34 architecture and outperformed the EfficientNetB0 architecture (87.1%). Both RNNs showed higher fps (17.9 for RNN Amoh and 19.8 for Vanilla RNN) with comparable, but lower median test accuracy (86.8% and 85.8%) to those of CNNs (Fig. 3D).

In summary, we were able to show that NSA signals contained sufficient data features for airway symptom detection in combination with DNN techniques. All investigated DNNs were, however, too large for wearable deployment. We thus proceeded to optimize the network topology with the focus on CNNs next, given their superior accuracy in airway symptom classification.

### CNN topology optimization using an evolutionary algorithm

An efficient classifier is integral for mobile health wearable deployment. Here, we investigated how to optimize CNNs in a directed fashion to allow both fast and accurate classification by being wearable and deployable. Evolutionary algorithms (Fig. 4A) were used to optimize CNN topology using either of the following two objective functions. Objective 1 (O1) was to maximize the validation accuracy of a CNN topology. Objective 2 (O2) was to maximize both validation accuracy and model’s processing frames per second (fps). Both objective functions were found to increase their median and maximum fitness across generations (see Fig. 4B). Due to the evolutionary algorithm and its mutation and cross-over features, there was a gap between median and maximum possible fitness. The distribution of individual topology genetics across generations are shown in Supplementary Fig. 4.

**Fig. 4.**
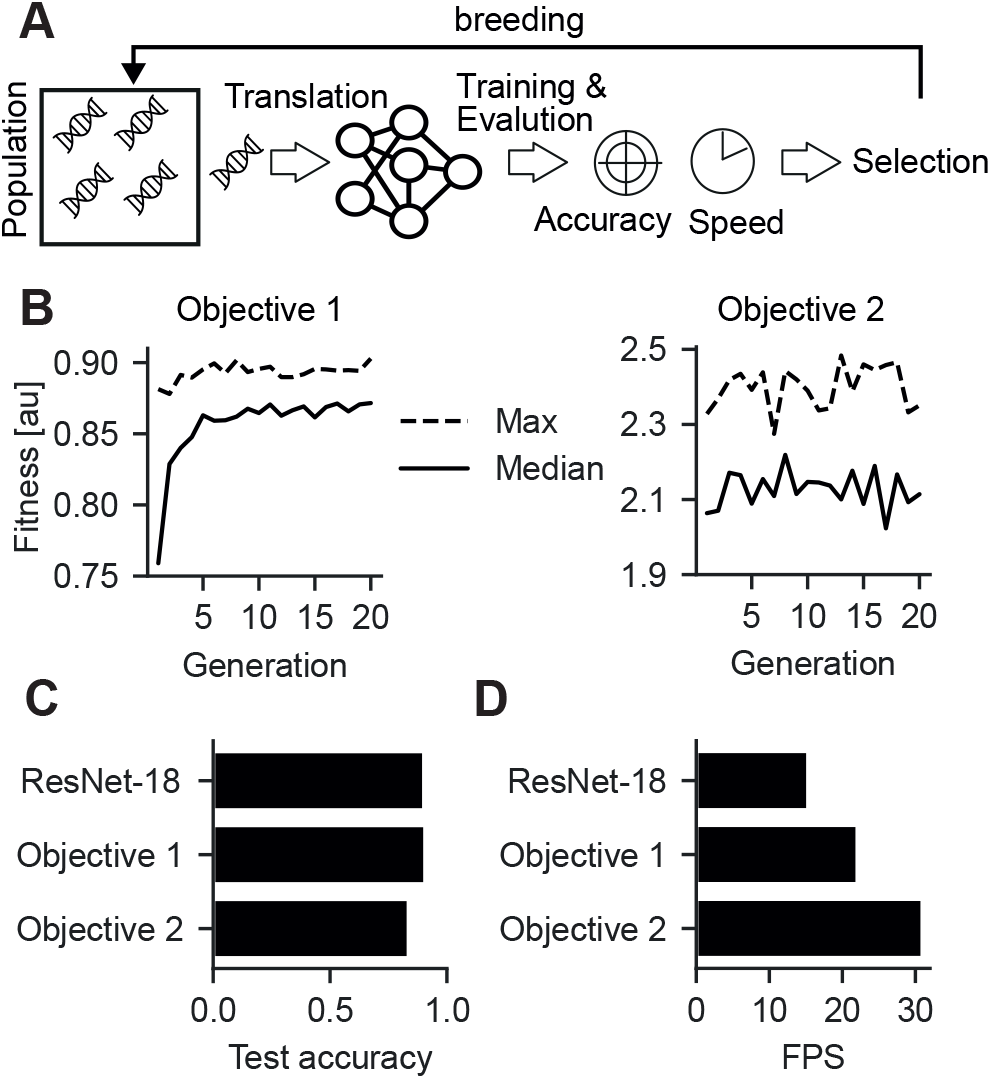
Evolutionary optimized deep neural networks are competitive with stateof-the-art networks. **A** We used a evolutionary approach to evolve neural network topology. We selected the network parameters from a genetic pool (Tab. 1), translated the parameters in a trainable deep neural network, trained and evaluated each individual and selected for the fittest. These were used to breed the next generation, employing cross-overs and random mutations. **B** Convergence of fitness across generations for objective 1, i.e. optimizing for accuracy only (left panel) and objective 2, i.e. optimizing for accuracy and inference speed (right panel). **C** Comparison of test accuracies from objective 1 and 2 models compared to baseline (ResNet-18). **D** Comparison of inference speeds of objective 1 and 2 models compared to baseline (ResNet-18).

In comparing with the baseline non-deployable CNN ResNet-18, the test accuracies of O1 and O2 were equally good or slightly worse (90.28% vs 90.75% (O1) and 83.67% (O2)) for one data split (Fig. 4C). However, the evolutionary algorithm boosted the processing speed from 15.3 to 31.0 fps (Fig. 4D, Tab. 2). The O2 model was faster than ResNet-18 and O1, confirming the fitness design and that it was possible to trade accuracy for inference speed (Table 2). The final O2 architecture consisted of 7,692 trainable parameters which was 0.069% compared to the baseline architecture (11,186,692 trainable parameters) and had a memory footprint of less than 150 kB. The determined network topology of the O2 model is shown in Supplementary Fig. 5.

**Table 2.** Evolutionary algorithm results compared to the baseline model.

| Architecture | Test Accuracy | FPS | Number of parameters |
| --- | --- | --- | --- |
| ResNet-18 (Baseline) | 0.903 | 15.3 | 11,186,692 |
| Objective 1 | 0.908 | 22.1 | 289,988 |
| Objective 2 | 0.837 | 31.0 | 7,692 |

To test performance in a real-world microcontroller environment, we deployed the ResNet-18 and the optimized O1 and O2 architectures to a developmental Deep Learning-enhanced board (EdgeBadge Board, see also Methods). Unfortunately, due to the large model sizes (44 and 3.5 MBs for ResNet-18 and O1, respectively), we were not able to deploy these models to the microcontroller, which was restricted to 512 kB of memory. However, once we converted and deployed our evolutionary optimized model O2 with TensorFlow Lite, we were able to gain 3.5 fps, which is considered a fair result for a non-optimized hardware board.

### Adaptability of evolutionary optimized CNNs to new data

Model fine tuning and transfer learning were used to evaluate how well the evolutionary optimized CNN (O2) could adapt to new data (Fig. 5A). The test accuracy of the Rainbow Passage dataset was preserved when fine-tuning the model with a combination of the Rainbow Passage dataset and a subset of the Vocal Stress dataset (9). However, the test accuracy of the new dataset converges at about 0.7, suggesting that the model was only able to fit the new dataset to a certain extent. When the Vocal Stress dataset was used in the training process (transfer learning) only, the pre-trained model was able to learn the representation quickly. While performing better on the test set, the test accuracy of the Rainbow Passage dataset dropped from 80% to 22.2%. Although the proposed architecture was specifically tuned to the Rainbow Passage dataset, it was capable of quickly learning new representations through transfer learning and fine tuning.

**Fig. 5.**
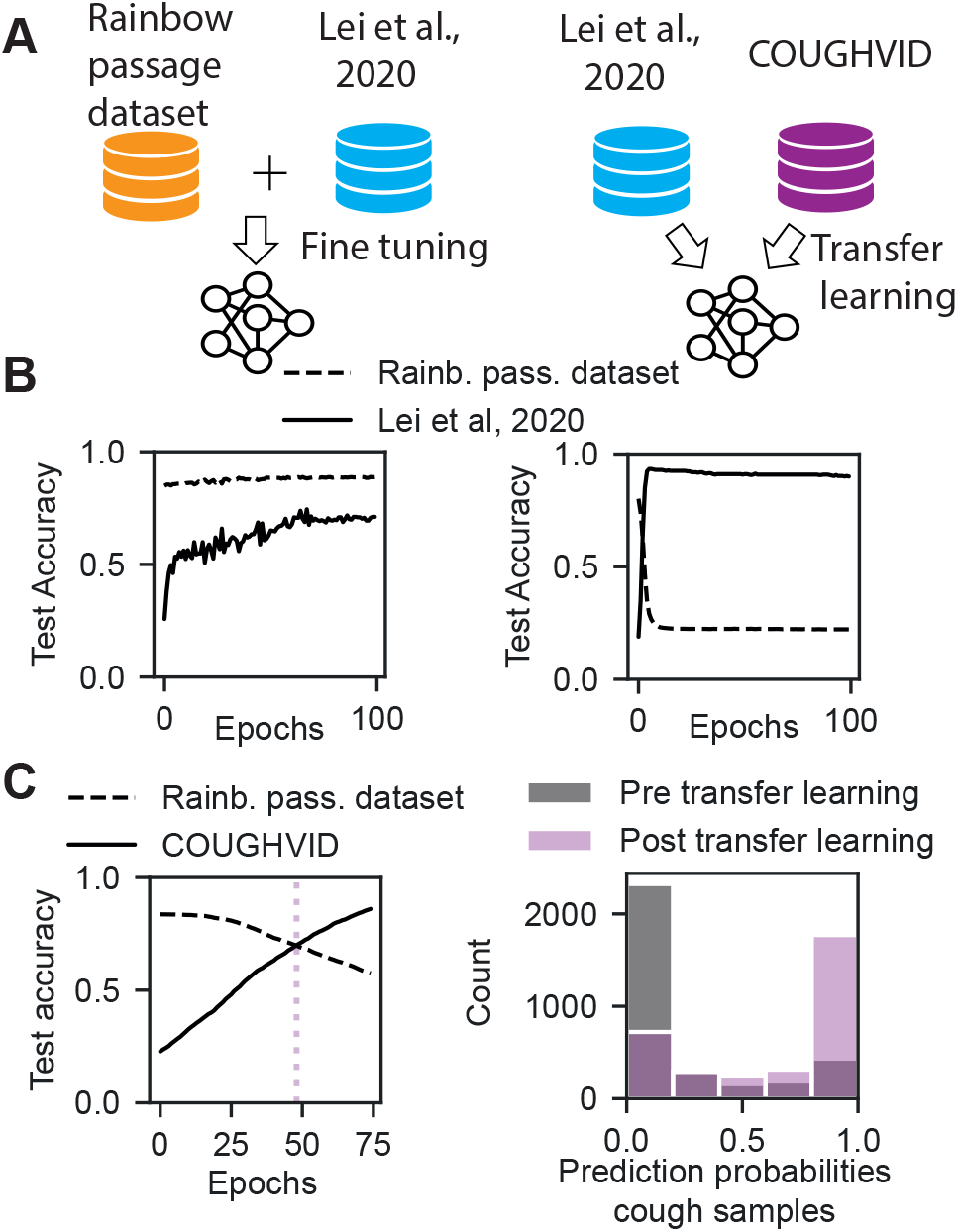
Fine tuning and transfer learning determined genetic algorithm model using two additional datasets. **A** Improving determined genetic algorithm architecture performance and robustness through fine tuning and transfer learning using related unknown datasets. **B** Evolution of test accuracy over epochs when using the dataset from Lei et al., 2020 with (left) and without (right) the Rainbow Passage dataset (Rainb. pass. dataset) for fine tuning and transfer learning the determined genetic algorithm model. **C** Transfer learning of the determined genetic algorithm architecture using the COUGHVID dataset.

The COUGHVID dataset contains a variety of cough audio samples gained from a crowdsourcing effort. To convert the audio samples to NSA space, a crucial step for testing the O2 CNN, we trained a U-Net-like architecture with the paired audio/NSA data extracted from the Rainbow Passage dataset by minimizing the mean squared error across spectrograms (Supplementary Fig. 6 for the workflow). We analyzed the conversion quality using the structural similarity index measure (SSIM, (39)). Our conversion procedure was able to produce valid spectrograms in NSA space. SSIM results showed that converted NSA spectograms were closer to real NSA spectrograms (mean SSIM=58.1 %) compared to audio-derived spectrograms (mean SSIM=37.6 %).

**Fig. 6.**
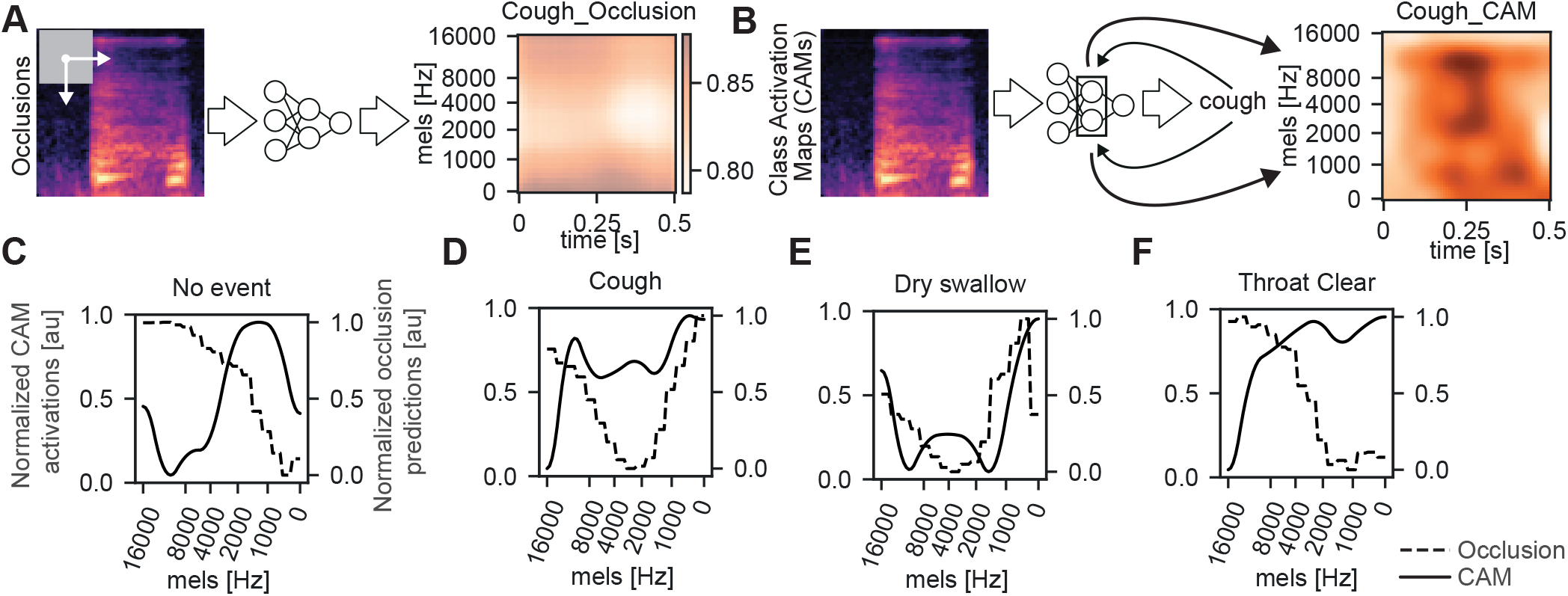
Specific frequency bands are important for cough detection. **A** Parts of the mel-spectrogram are occluded to determine which frequencies are important for successful event prediction, here cough. **B** Class activation maps are used to determine important spatiotemporal frequencies for event predictions, here cough. **C-F** Normalized class activation map (CAM, solid line) and normalized occlusion prediction (dashed line) profiles for no event, cough, dry swallow and throat clear, respectively.

We next analyzed the O2 model to classify the obtained NSA samples of the COUGHVID dataset. The prediction probabilities of cough samples were relatively low before transfer learning (Fig. 5C, gray bars). However, by fine-tuning the model on a few samples of previously unseen data (50 converted NSA cough samples from the COUGHVID database), the test accuracy on the remaining 3,138 cough samples increased dramatically from 22.8% to 70% on average (see Fig. 5C, pink bars). After 48 epochs, the model reached a similar performance for both datasets before overfitting was observed (Fig. 5C, left panel).

Although the O2 model might not be robust to different sources, the model was shown to quickly adapt to new datasets. This feature is particularly useful to support personalized wearable health technology. For cases like chronic airway diseases, an individual’s health data are dynamically evolved as functions of time history and personal profiles. The adaptability of the O2 model will allow self-learning and fine-tuning to improve its prediction accuracy when new data are supplied from individual patients.

### Airway symptoms cluster on specific frequency bands

Our next interest was to investigate if our optimized CNN O2 relied on specific frequency bands for airway symptom detection. That way, in case of confined frequencies, we would be able to further optimize the preprocessing steps and the DNNs to gain potentially even smaller or more robust models for mobile and wearable applications. We focused on two complementary approaches: occlusion experiments (38) and CAMs (36). The occlusion experiments (Fig. 6A, Supplementary Fig. 7) showed that mel-frequency bands up to 8000 Hz were most important for classifying coughs, dry swallows and throat clears (Fig. 6D,E,F). Mel-frequency bands of up to 2000 Hz were important for correct classification of no event (Fig. 6C). When analyzing the respective CAMs, we found confirming results for each event, with the exception of dry swallow (Figure 6C-F, Supplementary Fig. 8 and 9). Class activations were higher in the same mel-frequency bands where predictions dropped off when the bands were occluded. Taken together, airway symptom features were restricted to specific frequencies, allowing not only trustworthy applications but also leaner future models.

## Discussion

In this work, we showed that a scripted, tiny dataset was sufficient to train DNNs in the classification of airway symptoms on unseen data with satisfactory testing and validating accuracies. Using evolutionary algorithms, a neural topology was optimized for accuracy and inference speed that was less than 150 kB in size, which is on par with large state-of-the-art architectures. Such low size footprints are important for downstream application on mobile devices and wearables. Further, we found that specific frequency bands were important for airway symptom identification, which is essential for us to tailor the proposed DNNs in the future and to solidify trust in wearable health devices.

### Cough prediction

Coughing is a common symptom across multiple airway-related diseases such as asthma, COPD and COVID-19. Cough sounds serve as one common digital audio biomarker in mobile health technologies (15, 16, 31). For example, coughs have been used to predict COVID-19 positivity (40, 41). In this work, we used the COUGHVID database (21), a large crowdsourced database containing audio recordings with a wide range of perceptual audio quality of mainly cough sounds. A data cleaning strategy was tailored to extract cough sounds and convert the microphone audio signals to NSA space for model evaluation. Our proposed models were able to adapt to this diverse database. In contrast to other works, we specifically aimed for a deployment of our DNNs classification algorithms on low-power, computational restrictive wearable platforms (42, 43).

### Unboxing deep neural networks

Explainable AI is integral to advance and translate the technologies to clinical applications (44, 45). The occlusion experiments and CAMs (Fig. 6) identified frequencies that were specific to airway symptom prediction. As a sanity check, we confirmed that log-mel-spectograms classified as *no event*, which typically contained human speech signals, showed important fundamental and harmonic frequencies of up to 2000 Hz as expected in human speech. We also confirmed that CAMs were similar across architectures (such as ResNet-18 and the O2 optimized model), suggesting that the same concepts were learnt, despite the fact that the latter network features less than 1 % of the trainable parameters of the former. More recent technologies of network activation, such as DeepLift (46) and Grad-CAM (47) can also be included in future investigations.

### Limitations and shortcomings

Despite the fact that coughs were classified with satisfactory accuracies with our CNNs, the heterogeneity of the data resulted in a large fraction of false positives. The expansion of our current training set, i.e. the Rainbow Passage dataset, may help to further improve the classification accuracy and especially robustness to various sources. Nevertheless, with our limited training data, we already gained competitive results. We also noted that the genetic pool in our evolutionary algorithm was relatively limited and can be further expanded, for example with the use of depth-wise convolutions (48) or compound scaling (29). In future, we will investigate additional topology optimizations and test the resulting topologies in a real-world application. In this work, we achieved the first step of developing effective and explainable AI algorithms for long-term remote monitoring of airway symptoms by mechano-acoustic wearables.

## DATA AVAILABILITY

Data will be available upon reasonable request.

## CODE AVAILABILITY

The code for evolutionary algorithms, occlusions and class activation maps are found online at https://github.com/ankilab/AirwaySymptomDetection.

## ACKNOWLEDGEMENTS

We thank Tobias Kist for help with electronics. We acknowledge research grants from the Canadian Institutes of Health Research [PJT-156412], The Centre for Research on Brain, Language and Music Research (CRBLM) Incubator Awards (NLJ) and Canada Research Chair research stipend (NLJ). The CRBLM is funded by the Government of Quebec via the Fonds de Recherche Nature et Technologies and Société et Culture. The presented content is solely the responsibility of the authors and does not necessarily represent the official views of the above funding agencies.

## AUTHOR CONTRIBUTIONS

NLJ and AMK conceived and supervised the project. ZL designed the NSA board. LM and RG annotated data. RG analyzed the data, trained and evaluated deep neural networks, and conceived evolutionary algorithms. RG, AMK and NLJ wrote the paper. NLJ and AMK revised the paper critically for important intellectual content.

## Supplementary Figures

9 Supplementary Figures

## Supplementary Material

1 Supplementary Material: Rainbow passage script

