## Supplementary Figures for "Efficient and Explainable Deep Neural Networks for Airway Symptom Detection in Support of Wearable Health Technology"

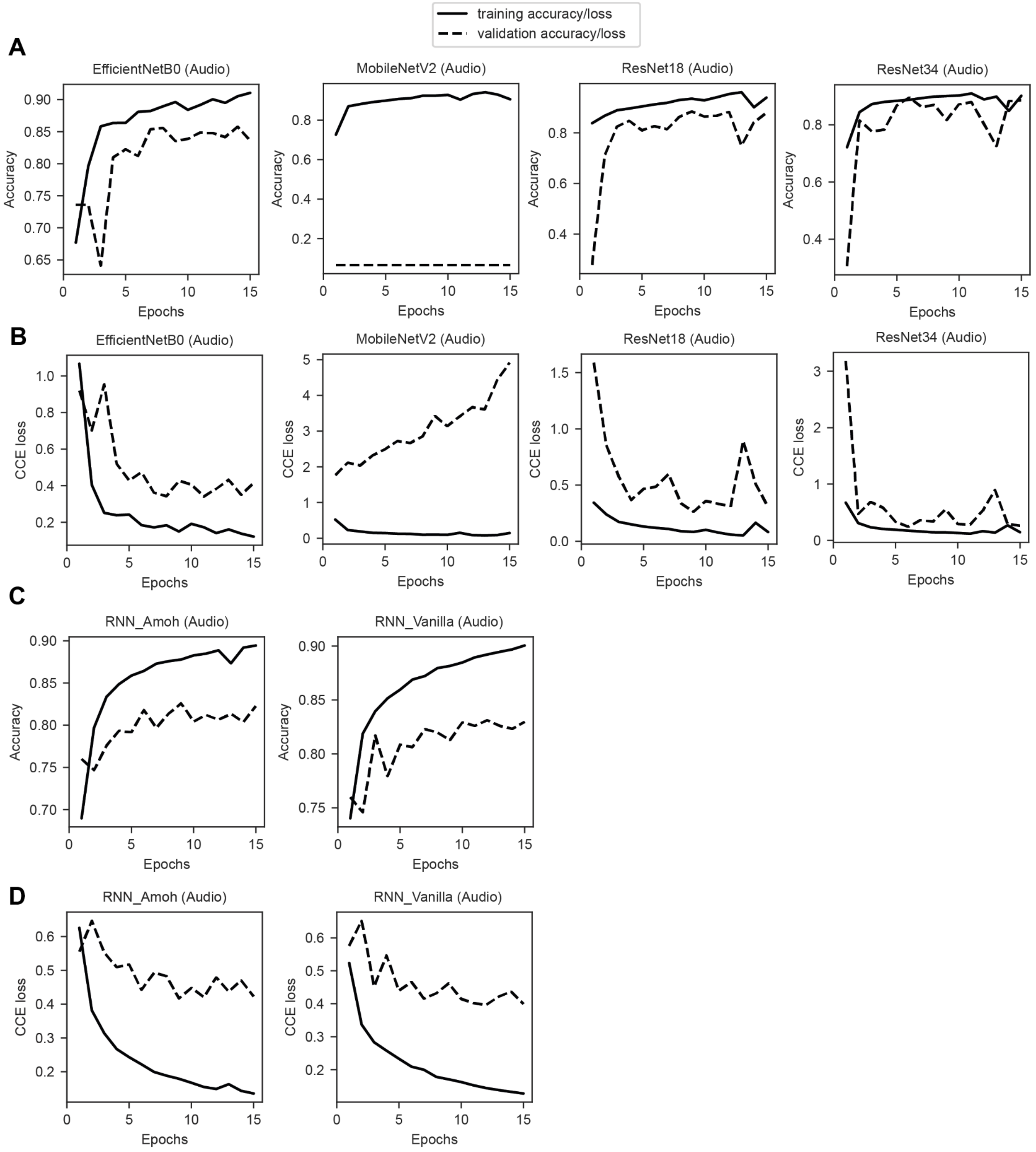


**Supplementary Figure 1. Convergence of Deep Neural Networks on audio data. A.** Accuracy for several CNN architectures. **B.** Categorical crossentropy loss for several CNN architectures. **C.** Accuracy for two RNN architectures. **D.** Categorical crossentropy loss for two RNN architectures.

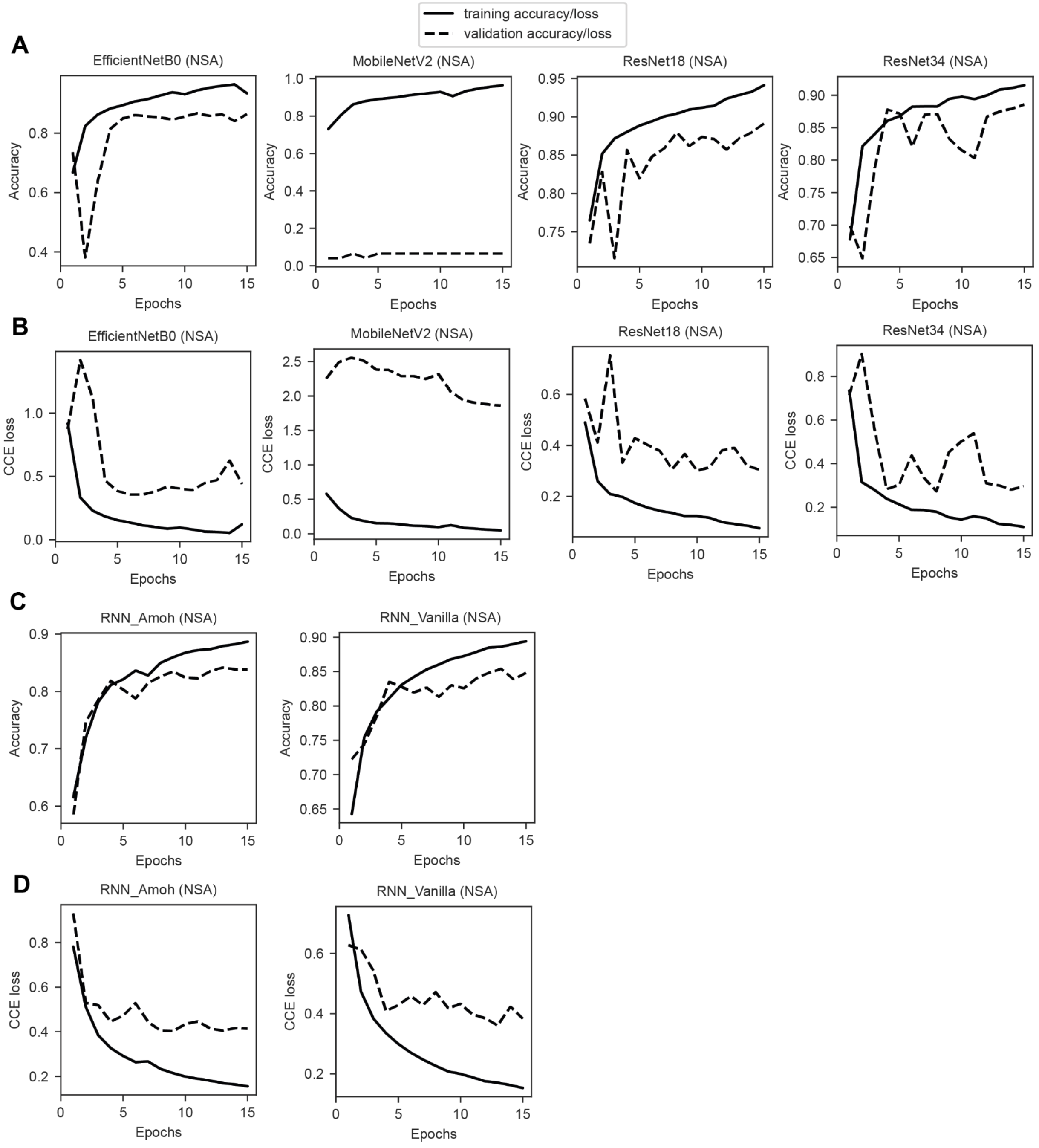


**Supplementary Figure 2. Convergence of Deep Neural Networks on neck-surface accelerometer (NSA) data. A.** Accuracy for several CNN architectures. **B.** Categorical crossentropy loss for several CNN architectures. **C.** Accuracy for two RNN architectures. **D.** Categorical crossentropy loss for two RNN architectures.

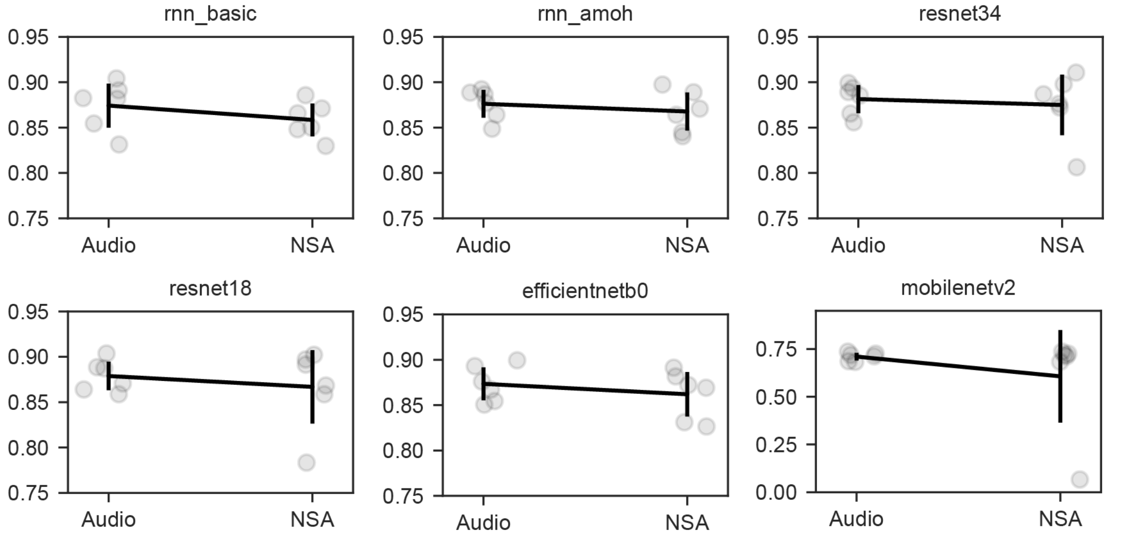


**Supplementary Figure 3. Comparison of audio and NSA data for each deep neural network.** From left to right, top to bottom: Vanilla RNN, RNN Amoh et al., ResNet-34, ResNet-18, EfficientNetB0, MobileNetV2.

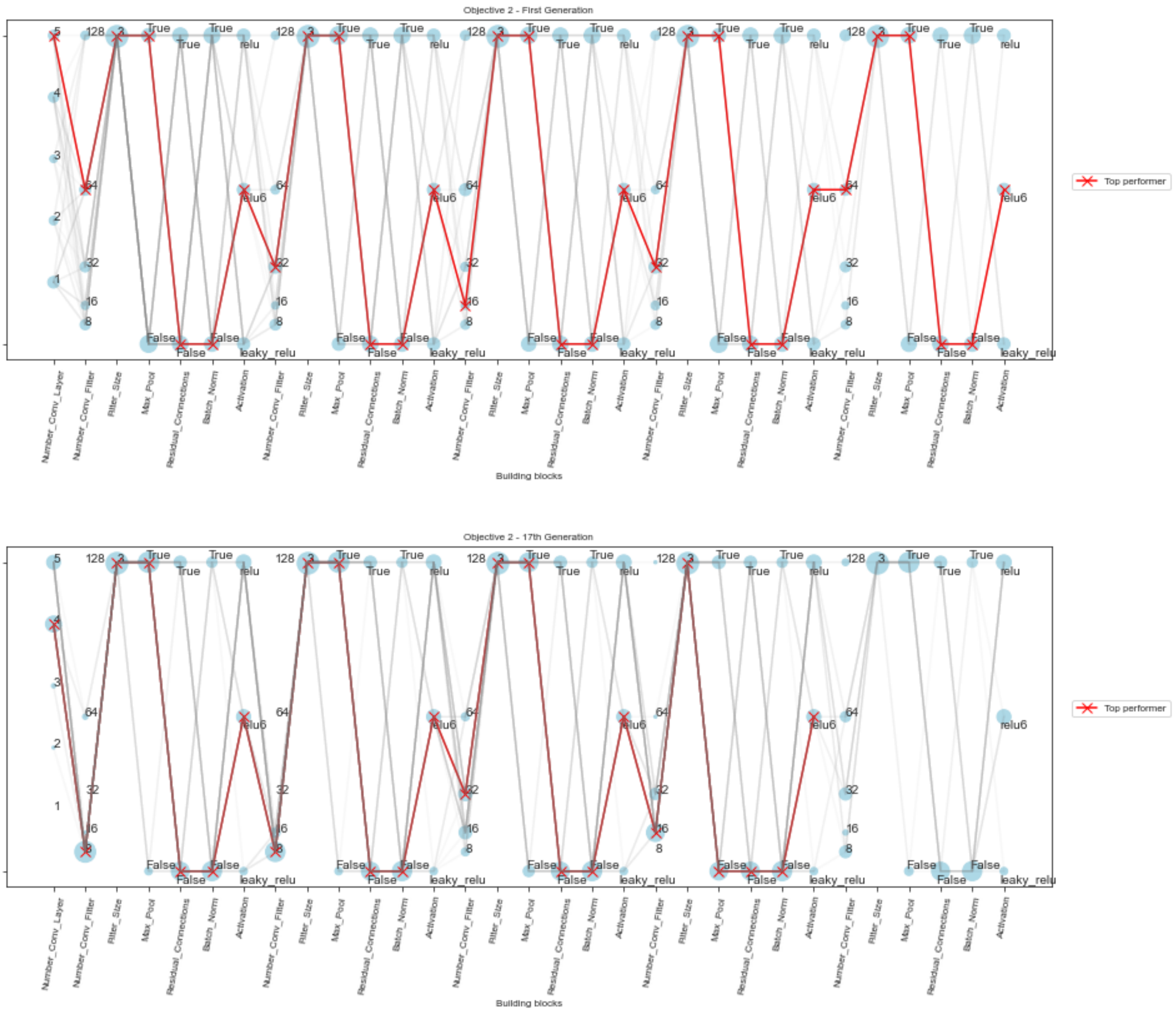


**Supplementary Figure 4. Distribution of network topology genetics across generations for Objective 2.**

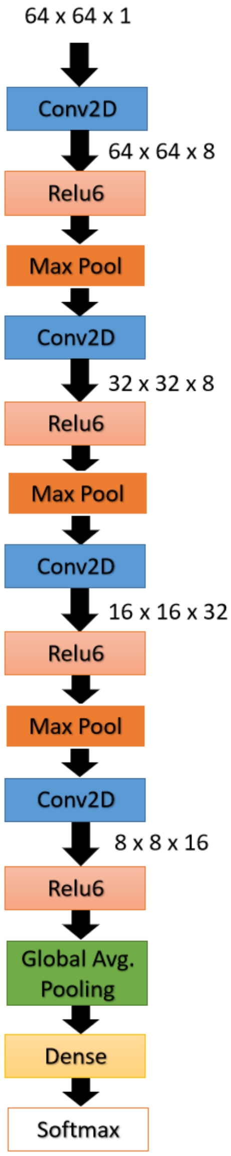


**Supplementary Figure 5. Network topology determined by genetic algorithm for Objective 2.**

**A.**

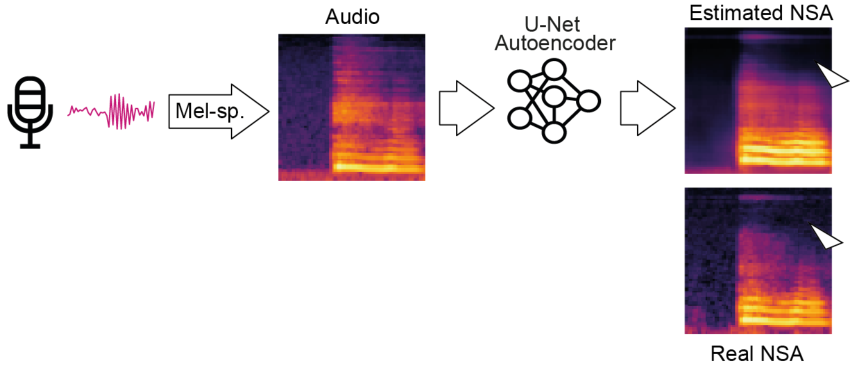


**B.**

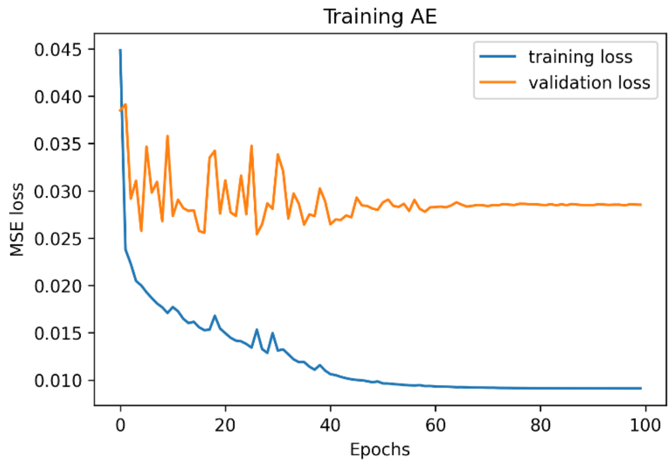


**Supplementary Figure 6. NSA data generation using a U-Net architecture. A.** Overview of audio-to-NSA data conversion and exemplary converted NSA spectrum. Note the loss of high frequencies. **B.** Convergence of deep neural network across epochs.

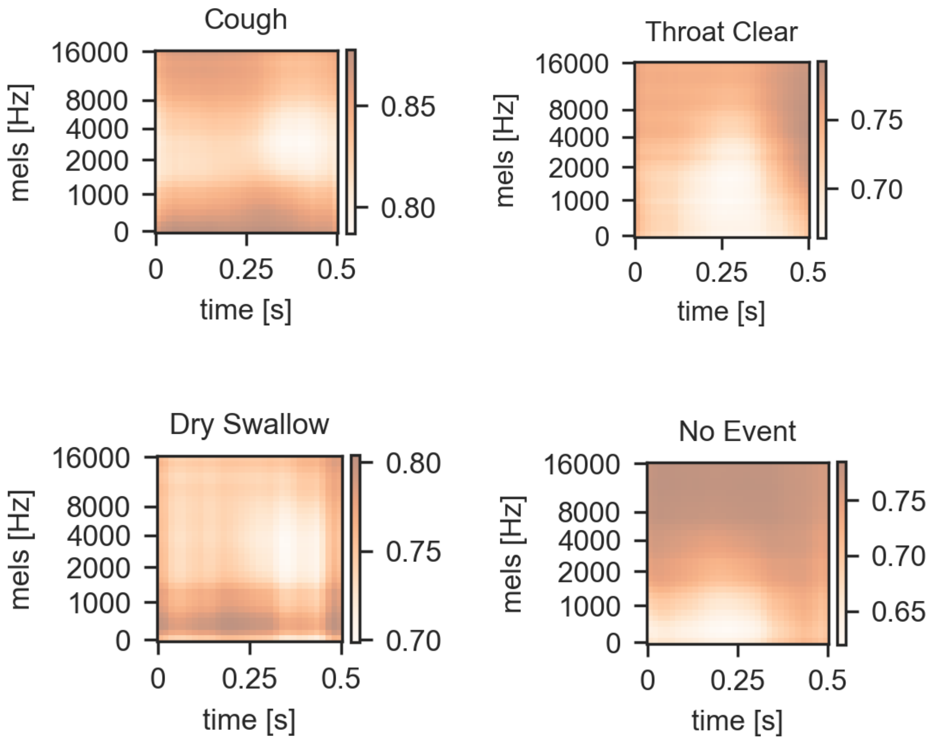


**Supplementary Figure 7. Occlusion experiments for airway-related symptoms.**

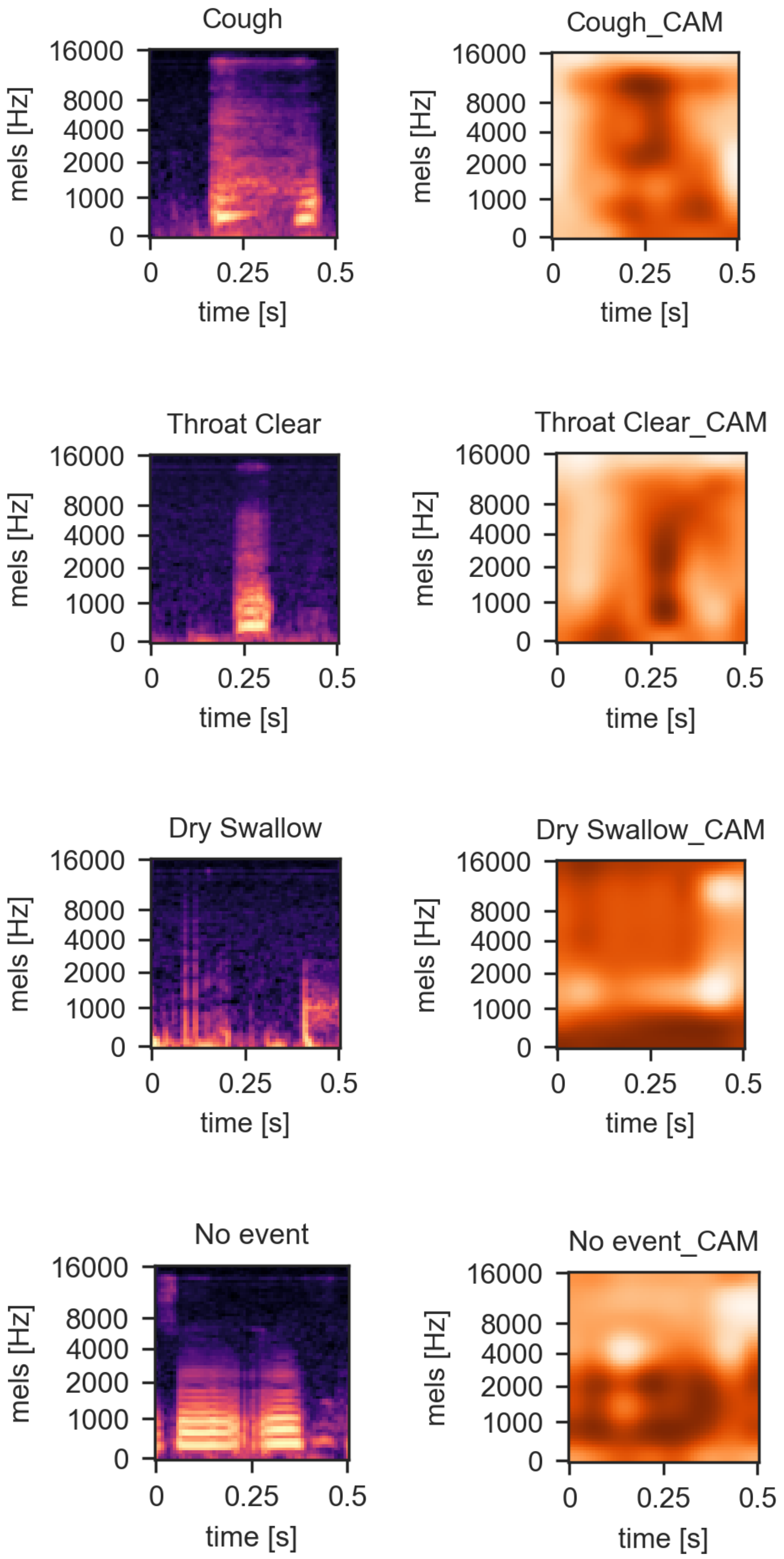


**Supplementary Figure 8. Examples of class-activation maps (CAMs, right panels) for airway-related symptoms (spectrograms left panels).** Darker areas result in higher class activations.

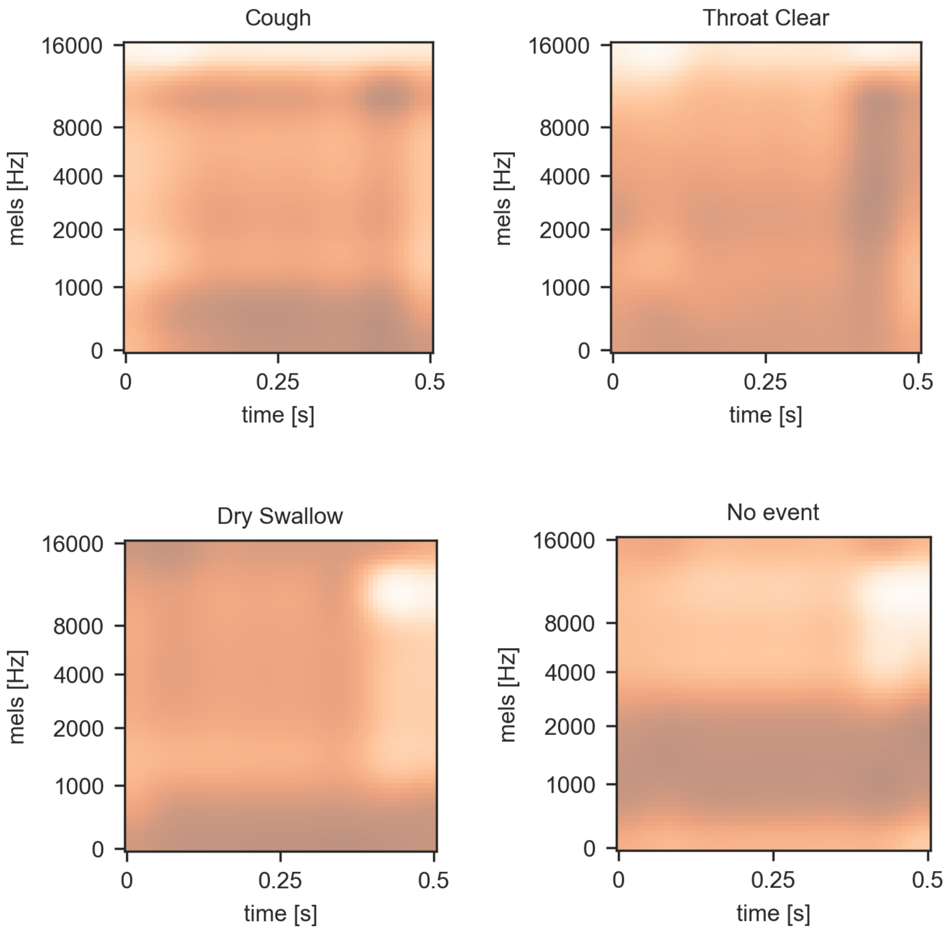


**Supplementary Figure 9. Average class-activation maps (CAMs) for airway-related symptoms.** Darker areas result in higher class activations.
