## Supplementary Material for "Efficient and Explainable Deep Neural Networks for Airway Symptom Detection in Support of Wearable Health Technology"

SCRIPT
"Rainbow Passage"

***(Cough)*** x1

***(Swallow)*** x1

***(Throat Clear)*** x1

When the sunlight strikes ***(cough)*** raindrops in the air, they act as a prism and form a rainbow ***(Throat clear)***. ***(Swallow)*** The rainbow is a division of white light into many beautiful colors ***(throat clear).*** These take the shape of a long round arch ***(cough),*** with its path high above ***(swallow),*** and its two ends apparently beyond the horizon. There is, according to legend, a boiling pot of gold at one end ***(throat clear)***. People look ***(cough)***, but no one ever finds it. When a man looks for something beyond his reach ***(swallow),*** his friends say he is looking for the pot of gold at the end of the rainbow. ***(Throat clear)*** Throughout the centuries people have explained the rainbow in various ways ***(cough)***. Some have accepted ***(cough)*** it as a miracle without physical explanation. To the Hebrews ***(cough)*** it was a token that there would be no more universal floods ***(swallow)***. The Greeks used to imagine that it was a sign from the gods ***(swallow)*** to foretell war or heavy rain. The Norsemen ***(throat clear)*** considered the rainbow as a bridge over which the gods passed from earth to their home in the sky ***(throat clear)***. Others have tried to explain the phenomenon physically ***(cough)***. Aristotle thought that the rainbow was caused by reflection of the sun's rays by the rain ***(cough)***. Since then ***(swallow)*** physicists have found that it is not reflection ***(throat clear)***, but refraction by the raindrops which causes the rainbows. Many complicated ideas about the rainbow have been formed ***(swallow)***. The difference in the rainbow depends considerably upon the size of the drops ***(throat clear)***, and the width of the coloured band increases as the size of the drops increases. The actual primary rainbow observed ***(cough)*** is said to be the effect of super-imposition of a number of bows. ***(Cough)*** If the red of the second bow falls upon the green of the first ***(throat clear)***, the result is to give a bow with an abnormally wide yellow band ***(swallow)***, since red and green light when mixed form yellow. ***(Swallow)***This is a very common type of bow ***(swallow)***, one showing mainly red and yellow, with little or no green or blue ***(throat clear)***.

***(Cough)*** x1

***(Swallow)*** x1

***(Throat Clear)*** x1
